# A Study of Coronary Artery Disease in the Left Circumflex Artery via Artificial Vessel Restoration

**DOI:** 10.1101/2020.05.27.119628

**Authors:** Akhil Paulraj, Fei Liu

## Abstract

Diagnosis, risk analysis, and treatment of coronary artery disease (CAD) can be improved with a better understanding of cardiovascular flows. Numerical simulations can assist in achieving this understanding. The objective of this study is to compare the dynamics of blood flow in a diseased left circumflex artery (LCX) and its artificially restored counterpart representing its healthy state. This comparison is made to identify flow characteristics in the restored vessel that contribute to the development of CAD. The diseased LCX geometry was derived from computed tomography angiography data. The stenosed region of the diseased LCX was repaired by manually redefining cross-sections of the vessel, creating the restored geometry. To account for inaccuracies, variations of the restored LCX were made by dilating the repaired surface. Numerical simulations were conducted on all geometries and the results were compared. Alongside expected low wall shear stress, a region of high vorticity was present in all of the restored vessels near the location where CAD developed in the diseased vessel. Therefore, this research suggests that flow vorticity is relevant in assessing the risk for CAD, potentially improving the accuracy of non-invasive, computational diagnosis. Such improvements can also help avoid unnecessary invasive diagnosis methods and minimize risk.

## 1 Introduction

Cardiovascular disease is the leading cause of death in the United States, reported by Benjamin et al. [2019] from the American Heart Association. Coronary artery disease (CAD) is the most common type of heart disease, and 6.7% of all adults in the United States over the age of 20 have CAD [Benjamin et al., 2019]. To garner more knowledge about cardiovascular disease and specifically CAD, computational methods have been employed as a research tool to study the fluid dynamics of cardiovascular flows.

Research on cardiovascular flows began with Leonardo da Vinci in the 15th century, who worked on in vivo studies with pigs to examine the flow of blood. da Vinci also constructed physical glass models of the heart and drew comparisons with the dynamics of water; in this manner, he recognized the importance of hemodynamics when studying cardiovascular flow [Sterpetti, 2019]. The basis of modern views of the cardiovascular system is rooted in the work of William Harvey, who was the first to both recognize and document his discovery of blood circulation [Parker, 2009]. Research on atherosclerosis, and in turn CAD, truly began only in the 19th century, as researchers began to hypothesize about the importance of wall shear stress (WSS). For many years, there existed a controversy between two contradicting hypotheses. One explanation argued that high WSS favors atherogenesis due to the mechanical damage caused by high WSS in the endothelium. The other explanation argued that low WSS favors atherogenesis because it is associated with arterial wall mass transport [Dhawan et al., 2010]. It has since been established that the second hypothesis is correct, also implying the significance of low WSS in the development of CAD [see Heo et al., 2014, Chiu and Chien, 2011, Nigro et al., 2011]. Disturbed blood flow in arterial bifurcations, branch ostia, and vessel curvature have also been shown to be prone to atherosclerosis [Nigro et al., 2011].

A more complete understanding of the hemodynamics contributing to the development of CAD is needed to improve upon existing risk analysis and diagnosis methods. By improving the accuracy of simulation-based methods, patients will have access to a cheaper, and often more accurate, way to be diagnosed.

The standard method for the diagnosis of patients with CAD is invasive coronary angiography (ICA) [Hulten and Carlson, 2013]. In ICA, a catheter is inserted into an artery, a dye is injected through the catheter into the coronary arteries, and X-Ray images are taken to inspect the vessels in question. Despite being the current standard for diagnosis, ICA has many drawbacks; for example, there may be complications that result from the invasive nature of ICA including site bleeding, deterioration of renal function, cerebral embolism and allergic reactions [Borren et al., 2015]. In addition, ICA is often unnecessary altogether: multiple registries have recognized that ICA is a low-yield diagnostic test. For example, analysis of the National Cardiovascular Data Registry CathPCI Registry done by Sahni and Tobis [2018] showed that only 40% of patients who underwent ICA had obstructive CAD. Moreover, the cost of ICA in comparison to simulation based methods that only require computed tomography angiography (CTA) is significantly higher [Darlington et al., 2015]. Hence, the unnecessary usage of ICA is needlessly expensive, which can be avoided by a simulation-based approach. For these reasons, ICA is only suitable for high-risk patients. Furthermore, the accuracy of ICA is limited: ICA tends to either underestimate or overestimate the severity of a stenosis [Borren et al., 2015, Skelly et al., 2016]. It should be noted that simulation based analysis methods still require data from CTA, which is not a completely non-invasive process, in that a dye must still be injected into the body through a vein in the arm; however, it is a far less invasive and expensive procedure in comparison to an ICA, which involves a catheder as aforementioned. Currently, simulation based diagnosis via CTA analysis has comparable accuracy to that of ICA, and with improved understanding of the hemodynamics associated with CAD, simulation based diagnosis will become more accurate [Gorenoi et al., 2012].

The most popular non-invasive method for CAD diagnosis is the cardiac stress test [Kharabsheh et al., 2006]. The use of an electrocardiogram in tandem with a cardiac stress test is a common, non invasive method to diagnose CAD. However, occlusions, particularly in the left circumflex artery (LCX), are under diagnosed using this method [White et al., 2017]. Moreover, patients may be unable to perform exercise as part of the stress test. There are also several other factors that may result in poor electrocardiogram readings; for example, medications such as digoxin can result in false positive S-T segment fluctuations, while other abnormalities can result in uninterpretable results [Skelly et al., 2016]. Such issues can have serious implications, and lead to unnecessary ICA which both introduces risk and is expensive.

LCX occlusions are responsible for 1 in every 5 myocardial infarctions [White et al., 2017]. Additionally, LCX occlusions have been found to present increased risk of heart failure and mortality in comparison to occlusions in the right coronary arteries and the left anterior descending arteries [Kozuch et al., 2015]. Given both the frequency and risks of CAD, it is clear that improving upon diagnosis methods is valuable. Therefore, due to the particularly high risk of LCX occlusions, an analysis of the hemodynamics associated with CAD in the LCX was conducted.

The purpose of this study is to compare the dynamics of blood flow in a diseased LCX and an artificial representation of its formerly healthy state, which is defined as the restored LCX. The characteristics of flow unique to the restored LCX are potentially associated with the development of CAD. Therefore, this comparison is made in order to identify cardiovascular flow characteristics that contribute to the development of CAD. It is expected that one of the identified flow characteristics is a region of low WSS, since it is established that low WSS contributes to the formation of atherosclerotic lesions. Other identified characteristics can serve as indicators for high risk of CAD. Hence, the identified characteristics in this study are significant in improving the accuracy of non-invasive, computational risk analysis and diagnosis methods for CAD; if one of the identified characteristics is observed in a patient, it can be used as a metric supporting potential growth of CAD. Furthermore, the findings of this analysis are relevant to the development of CAD in the LCX as well as other blood vessels.

The restoration technique used in this study was inspired by Berg et al. [2019], who utilized Voronoi diagram based reconstruction and mesh sweep reconstruction to artificially remove stenosis in the renal arteries. In this study and others, a restoration procedure to artificially create a healthy vessel is used largely because it is both difficult and rare to have access to data of a patient before and after atherosclerotic stenosis. A potentially viable solution to this problem is to artificially repair the vessel geometry as done in this study.

The principle of artificially restoring a diseased vessel in order to conduct a hemodynamic study was also implemented by Li et al. [2018] in “Retrospective Study of Hemodynamic Changes Before and After Carotid Stenosis Formation by Vessel Surface Repairing”, where an analysis of the hemodynamics associated with stenosis in the carotid artery was conducted. The carotid sinus was reconstructed and compared to the diseased vessel to gain further insight into the hemodynamics associated with atherosclerotic stenosis.

## 2 Materials and Methods

### 2.1 Data

The vessel geometry used for simulation is derived from the volumetric CTA data of a diseased LCX obtained from the Cardiovascular and Pulmonary Model Repository, constructed under Contract HHSH268201100035C from the National Institutes of Health under the direction of the National Heart, Lung, and Blood Institute Corporation [2009]. This data is not open source; however, it can be readily purchased from the aforementioned repository, where it is labelled as “OSMSC0158”.

The CTA data encompasses a square-prism region with a base of 19.93 cm and a height of 12.6 cm; hence, the data covers a volume of 5.00 × 10^3^ cm^3^.

The patient is a 65 year old male who is 64.3 inches tall and weighs 108.7 lbs. The patient’s blood pressure is determined to be 131/68 mmHg, and the patient’s cardiac output is determined to be 4.21 L/min [Ellwein-Fix et al., 2011]. The patient has CAD in the LCX, and underwent percutaneous coronary intervention. The obtained CTA data was taken preoperatively. The CTA data was processed and analyzed in SimVascular, using a framework based on the Medical Imaging Interaction Toolkit. This framework allows the user to parse volumetric CTA data into the axial, sagittal, and coronal planes. This simplifies the process of defining the geoemetry of the LCX.

Data availability is a crucial factor in this area of research. If abundant datasets are made publicly available, it will enable large scale studies that can better validate hemodynamic patterns associated with CAD. The procedure in this study and others can be reiterated to identify the accuracy and tolerance of identified patterns in slightly different geometries. This will significantly improve the value of these studies as well as other approaches involving different techniques such as machine learning. Hence, limitations in data availability due to either privacy or commercialization greatly hinder progress in this field of research.

### 2.2 Theory of Simulation

The primary goal of this research is to study the flow of blood in a diseased and restored LCX. To conduct this comparison, computational fluid dynamics techniques are employed using the open source cardiovascular simulation tool, SimVascular, developed jointly by researchers at Stanford University and University of California, Berkeley [Updegrove et al., 2017]. SimVascular version 2019.07.10 was used.

To model blood flow, SimVascular solves the incompressible Navier-Stokes Equations,

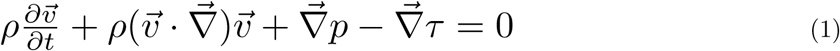

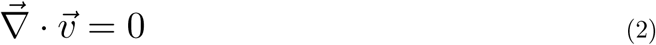

where *ρ* is blood density, 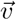 is the the velocity of the blood flow, *p* is the pressure, and *τ* is the viscous portion of the stress tensor [Updegrove et al., 2017]. These are the standard Navier-Stokes equations solved in computational fluid dynamics. The numerical solver in SimVascular uses a finite element discretization based on an unstructured mesh to obtain numerical solutions to the Navier-Stokes equations.

### 2.3 Simulation Setup for Diseased LCX

To define the computational domain for simulation, a path for the diseased LCX must first be defined. In SimVascular, a path is defined by two sets of points: control points, which are selected by the user, and path points that subdivide control points in regular intervals. Control points are chosen by tracing the vessel in the volumetric CTA data, which can be done by slicing the image along the axial planes. The approximate total number of path points is defined by the user. The two sets of points combined define a vessel path, where the path is defined by an interpolating spline. To define the LCX path, 15 control points were defined, alongside a total of 106 path points.

To construct a model of the vessel, 2D segmentations representing the cross sections of the lumen are defined. SimVascular renders an image of the plane perpendicular to the vessel path at a given point along the path. The path can be traversed using the image resclice probe. To segment the diseased LCX, 19 segments approximating the vessel cross-sections as circular were used. However, in order to accurately capture the diseased region, three segments in the diseased region were manually defined using a spline polygon. These segments were traced over the cross-sections of the model obtained from the Cardiovascular and Pulmonary Model Repository, which was constructed in further detail using optical coherence tomography [Ellwein-Fix et al., 2011]. The set of segmentations are then lofted together with B-splines to create a solid model.

The solid model is then used to create a 3-D mesh of the computational domain, resulting in discretization suitable for the numerical solver in SimVascular. A tetrahedral mesh based on 3D Delaunay Triangulation [Si, 2015] is created for the LCX using SimVascular. The mesh has 160272 tetrahedral elements, 30513 nodes, 30315 edges, and 20210 faces. The global maximum for the tetrahedron edge size is 0.0313 cm.

### 2.4 Simulation Conditions and Properties

A steady flow simulation was conducted (as opposed to pulsatile or Womersley flow), as it has been shown to be sufficient to describe the hemodynamics of coronary blood flow in stenosed vessels [Mates et al., 1978]. The dynamic viscosity of blood was defined as 0.04 poise, and the density of blood was defined as 1.06 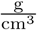. A no-slip boundary condition was implemented, and the walls of the vessel were considered to be rigid.

A velocity is prescribed to the inlet to serve as a boundary condition. A parabolic inlet velocity profile was used. To determine the inlet flow rate, the patient’s cardiac output was used. The cardiac output received by the LCX was calculated on the basis that the coronary arteries receive .05 of the total cardiac output, .84 of which is allocated to the left main coronary artery from which the LCX receives .33 [Ellwein-Fix et al., 2011]. The calculated values are shown in table 1.

**Table 1:** Inlet Boundary Condition Calculation

| Patient cardiac output | LCX fraction of output received | LCX output received |
| --- | --- | --- |
| $4.21 \frac{\text{L}}{\text{min}}$ | 0.01386 | $0.97251 \frac{\text{cc}}{\text{s}}$ |

Hence, the inlet flow rate is 0.97251 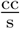.

A resistance is assigned to the vessel, serving as an outlet boundary condition by making the face outlet into a weakly-pressure face. In this manner, the vessel obeys a form of Ohm’s Law for advective processes:

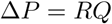

Δ*P* is the pressure drop across the vessel, which is analogous to voltage. *R* is the vessel resistance, which is analogous to the equivalent resistance of a circuit. *Q* is the volumetric flow rate, or received cardiac output, which is equivalent to current.

If the total peripheral resistance of blood flow in the patient is known, the ratio of the total cardiac output to the received cardiac output by the LCX can be used to determine the resistance of the LCX to be around 121953 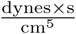.

To determine the time step size Δ*t* for the simulation, the Courant–Friedrichs–Lewy condition (CFL) is used to make an estimation. The CFL condition provides a relationship between the flow velocity, a temporal discretization parameter (time step size) Δ*t*, and a mesh discretization parameter *h* [Updegrove et al., 2017]:

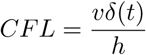

To obtain a reasonable time step size, the CFL number must be approximately 1.0 [Updegrove et al., 2017]. Using velocity calculated from the volumetric flow rate and the global maximum of the mesh element edge size for the mesh discretization parameter, the time step size is calculated to be 0.002 seconds.

Another factor must be considered: the number of time steps must provide sufficient time for the flow to reach a steady state. 500 time steps yields a physical time of 1 second, which is sufficiently long. The data is saved every 25 time steps.

### 2.5 Simulation Setup for Restored LCX and its Dilated Variations

The purpose of vessel restoration is to attempt to restore the diseased LCX to a geometry that accurately represents the healthy LCX before the development of CAD. This allows for an assessment of the hemodynamics in the FDR in order to determine potential dynamic factors that are associated with the development of CAD. In order to restore the diseased LCX, the three segments in the diseased region that were manually defined using a spline polygon to characterize the stenosis were removed, as shown in figure 1(b). These three segments were replaced by artificial, circular segments to represent a healthy vessel; this was done using a simplified version of mesh sweep reconstruction [Berg et al., 2019]. First, the centerline of the new vessel was computed and extracted as shown in figure 1(c). A linear function was used to determine the radius of circular segments that replace the diseased segments. These artificial segments are centered on the computed centerline, and have a radius determined by the function *r* = −.0035*x* + .288 based on the mesh sweep reconstruction algorithm, where *x* is the reslice value when traversing through the vessel path using the image resclice probe. The artificial segments are shown in figure 1(d).

**Figure 1:**
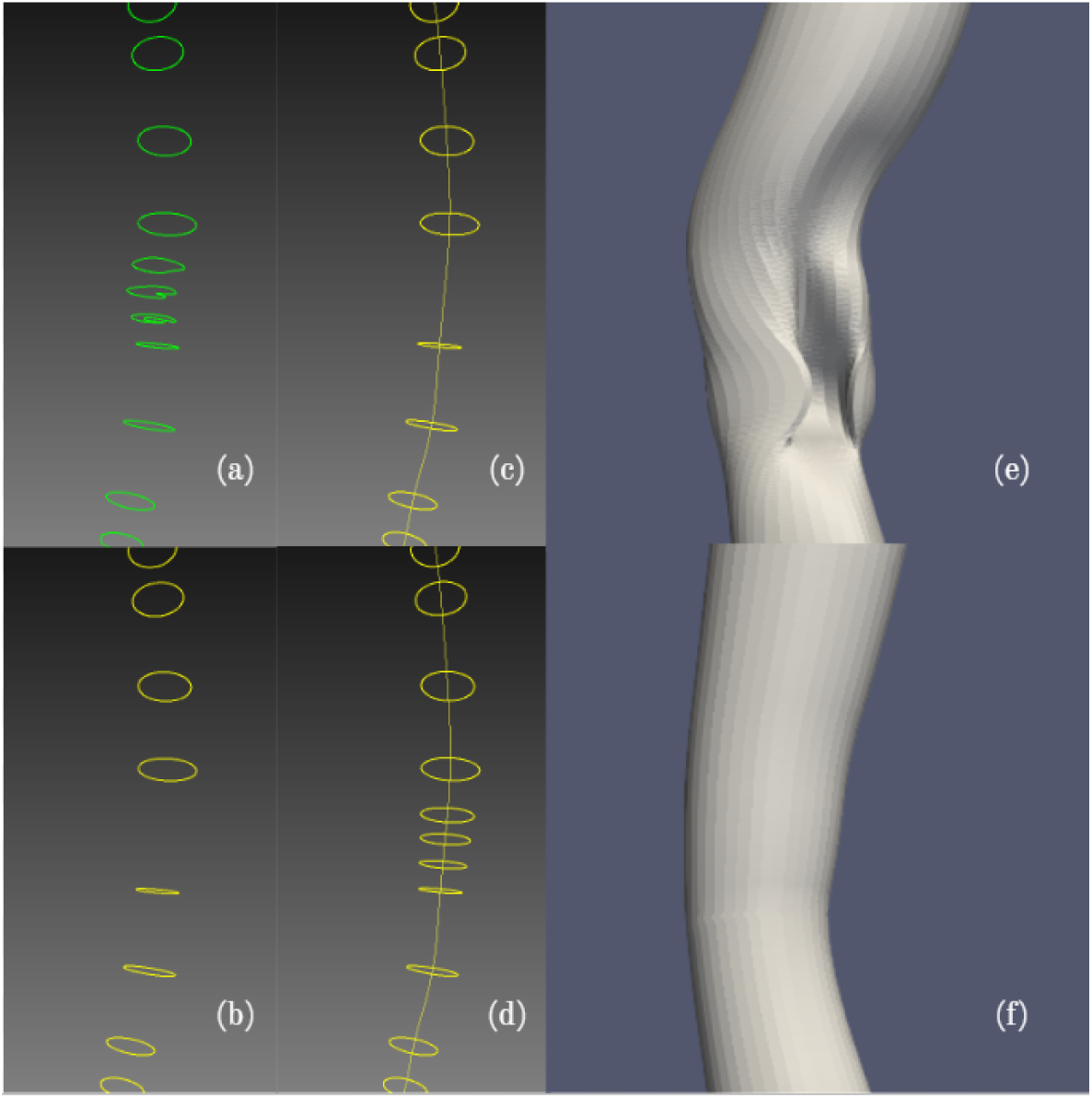
Outline of the diseased LCX Restoration Process. (**a**) Rendering of the segmentation of the diseased LCX. All segments are circular cross sections approximating the vessel geometry, except for the 3 diseased segments representing the stenosis. (**b**) The 3 diseased segments are removed as the first step in repairing the stenosis. (**c**) The centerline of the segmentation in (b) is computed and extracted. This is done in order to approximate the centerline of the initially healthy LCX. (**d**) 3 new segments are added to the segmentation to replace the 3 diseased segments from (a). The new segments are circular cross sections centered about the centerline from (c). The radii of these healthy segments is determined using a linear function such that the cross-sectional area changes gradually as shown. (**e**), the diseased LCX model, and (**f**), the restored LCX model, are also shown for comparison.

The new segmentation is lofted to create the restored vessel’s model. The diseased model is shown in comparison to the restored model in figure 1 parts (e) and (f) respectively.

A new mesh is created for the restored model, and simulations are run on this mesh using the same boundary conditions and general properties as the previous simulation.

The restoration technique used has its limitations in representing the initially healthy LCX of the patient. It is unlikely that the original, healthy geometry was smaller than the artificially restored geometry used in this study, simply because a further contracted geometry would intrude into the boundaries of the stenosed vessel. However, it is possible that the true vessel was larger than the artificially restored vessel used in this study. To account for this possibility and examine the tolerance of the hemodynamic characteristics identified in the artificially restored model, the three initially diseased segments in the LCX model which were then artificially replaced with circular segments were further dilated. This created three dilated variations of the restored LCX. Each of the three segments were dilated by a certain *δr* relative to its radius, *r*. Relative tolerances of 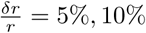,and 15% were tested, as shown in table 2. The hemodynamic traits of the dilated geometries were compared to that of the original restored geometry in order to validate the characteristics that were already identified.

**Table 2:** Segment Dilation Calculations

| Dilation Factor | Initial Segment Radius (cm) |  |  | Final Segment Radius (cm) |  |  |
| --- | --- | --- | --- | --- | --- | --- |
|  | Segment 1 | Segment 2 | Segment 3 | Segment 1 | Segment 2 | Segment 3 |
| 5% | 0.1270 | 0.1165 | 0.1095 | 0.1335 | 0.1223 | 0.1150 |
| 10% | 0.1270 | 0.1165 | 0.1095 | 0.1397 | 0.1282 | 0.1205 |
| 15% | 0.1270 | 0.1165 | 0.1095 | 0.1461 | 0.1340 | 0.1259 |

Simulation results were visualized using the open source post-processing tool ParaView[Ahrens et al., 2005, Ayachit, 2015]. ParaView version 5.7.0-RC1 was used. Quantities such as pressure and velocity were extracted from the simulation files and were exported to the ParaView tool. The resulting plots from ParaView consist of calculated quantities, such as WSS, over the unstructured tetrahedral mesh.

## 3 Results

The following graphs are volumetric, 3D plots of pressure, WSS, and vorticity. The corresponding color scales represent the values shown.

### 3.1 Comparison of Pressure Gradient in Diseased LCX and Restored LCX

In the pressure graph shown in figure 2(a), there is a pressure gradient ranging from 1.1×10^5^ Ba to 1.3×10^5^ Ba in the diseased LCX, which is a result of the assigned resistance boundary condition, *R*_LCX_. A noticeable change in pressure occurs near the diseased region, where high pressure is clearly seen directly before and above the diseased region due to the reduced cross sectional area from the stenosis that developed. Lower pressure is hence observed below the diseased region. This is expected, as the velocity of blood through the diseased region is increased according to the continuity equation due to decreased cross sectional area; therefore, a pressure reduction is reasonable considering Bernoulli’s principle. Moreover, the stenosis can be considered as an internal resistance of the vessel, which in turn must result in a pressure drop according to the equivalent Ohm’s Law for advective processes.

**Figure 2:**
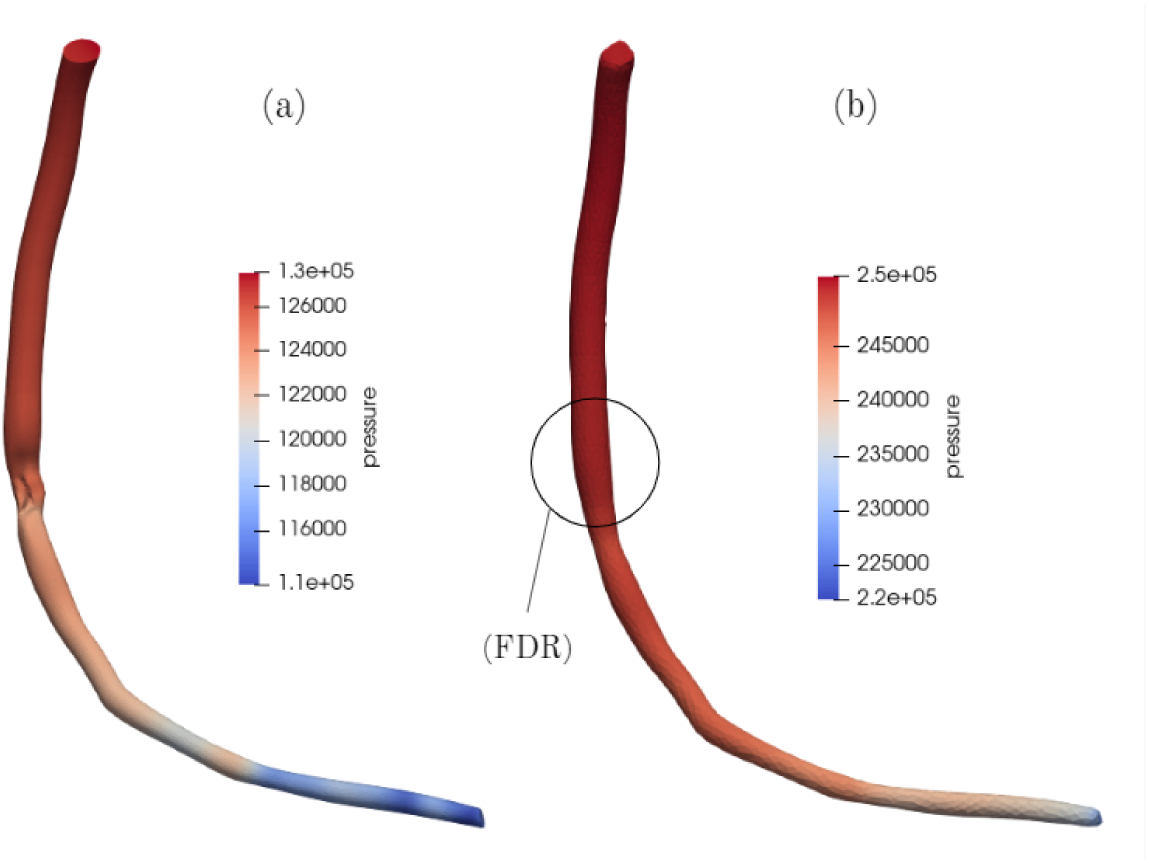
Pressure in the Diseased and Restored LCX, measured in Ba 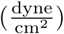. (**a**) Pressure in the diseased LCX. There is a pressure gradient from the inlet to the outlet, as indicated by the range of colors: the red at the inlet describes high pressure, and the blue at the outlet describes low pressure. There is an abrupt change in pressure at the diseased region due to the stenosis. This is reflected in the sharp color transition from red to light orange in the diseased region. (**b**) Pressure in the restored LCX. The formerly diseased region (FDR) is abbreviated and is circled. There is a gentler transition in color from the inlet to the outlet, as seen in the smooth color gradient from red to blue. This is clearly seen in the lack of a transition in color at the FDR.

Similar to the pressure graph of the diseased LCX, a pressure gradient exists in the restored vessel shown in figure 2(b) ranging from 2.2 × 10^5^ Ba to 2.5 × 10^5^ Ba. More importantly, the pressure gradient present in the restored vessel is smooth as there are no obstructions in the restored region, in contrast to the observed pressure drop at the diseased region in the diseased LCX. This is particularly clear when comparing the pressure change across the FDR in 2(b) to the diseased region in 2(a). This pressure characteristic is observed in the dilated variations of the restored LCX as well.

Hence, it is determined that noticeable pressure drops indicate obstruction of blood flow and possible stenosis. While identifying such pressure drops does not necessarily help with predicting the risk of CAD development, it can however identify regions where atherosclerosis has already begun to obstruct blood flow. This can in turn help prevent further growth of atherosclerotic lesions and its eventual repercussions such as myocardial infarction.

### 3.2 Comparison of Wall Shear Stress in Diseased LCX and Restored LCX

High WSS is observed in the diseased region, as shown in the shades of red featured in figure 3(a), reaching a maximum of 125.301 Ba. This local maximum results from the obstructive geometry of the stenosis. Therefore, the expected results are achieved, in that the presence of CAD is disrupting the flow of blood through the LCX.

**Figure 3:**
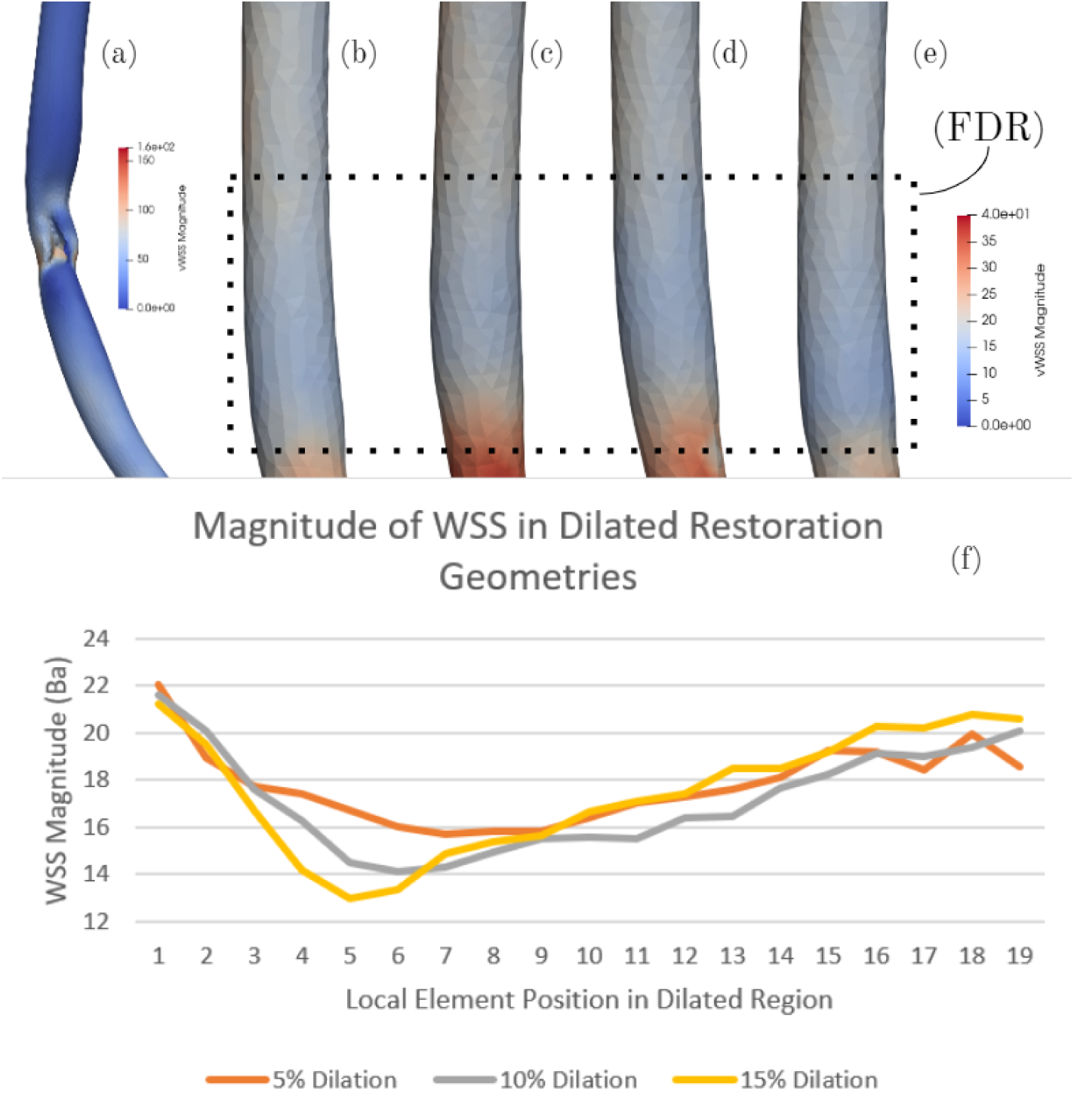
WSS in the diseased LCX, restored LCX, and dilated variations, Ba. The FDR is outlined by the dashed rectangle. (**a**) WSS in the diseased LCX. The blue shades throughout the vessel indicate shear stress that is weaker relative to the diseased region, which features relatively high wall shear stress as indicated by the shades of red and orange present. (**b**) WSS in the undilated restored LCX. Blood flow enters from the top of the image and exits through the bottom. Here, shades of blue indicate low WSS. In the figure, the FDR is dominated by low WSS and portrayed by the blue coloration. (**c**,**d**,**e**) WSS in the 5%, 10%, and 15% dilated variations of the restored LCX respectively. As shown, all three dilated variations feature low WSS in the FDR, indicated in blue. It is also noted that there are darker shades of blue present in the geometries that were more dilated. (**f**) Magnitude of WSS in dilated variations of the restored LCX. Each of the three dilated vessels were analyzed for WSS along its longitudinal axis. Elements were selected longitudinally across the FDR, such that the 0th local element position corresponds to an element downstream of the FDR and the 20th element corresponds to an element upstream of the FDR. In this manner, plots of WSS vs. local element position were created, illustrating that a local minimum of WSS is present in the FDR. The WSS of the 5%, 10%, and 15% dilated restored LCX is shown in orange, green, and yellow. The 15% curve achieves the lowest minimum WSS, and also reaches its minimum at the lowest element position (most downstream).

Upon restoration, the FDR of all the restored vessels feature low WSS as shown in figures 3(b,c,d,e). This contrasts with the high WSS in the diseased LCX in figure 3(a). In the undilated restored LCX, a local minimum in WSS of about 15.8 Ba is observed; this is significantly smaller than the high WSS of 125.301 Ba observed in the diseased LCX. This is in agreement with the fact that CAD eventually developed in this region, given that low WSS is a well-established contributor to atherosclerosis and the development of CAD. Therefore, the low WSS characteristic of the restored geometry supports the general validity and accuracy of the restoration process, as this is an expected characteristic of a region that developed CAD.

As aforementioned, there is a region of low WSS in the FDR of all three dilated variations of the restored LCX, as indicated in figures 3(c,d,e). This is in agreement with the results from the standard, undilated restoration simulation, which also featured a local minimum in WSS. Furthermore, the geometries with larger dilation factors have lower minimum values of WSS in the dilated region. The 15% dilated restoration geometry exhibits the lowest minimum WSS of 12.9665 Ba. The 10% dilated restoration geometry exhibits a minimum WSS of 14.1081 Ba. Meanwhile, the 5% dilated geometry exhibits the highest minimum WSS of 15.6945 Ba. Additionally, the minimum WSS in restoration geometries with larger dilation factors lies towards the downstream end of the dilated region. The minimum WSS in the 15% dilated geometry is the most downstream, whereas the minimum WSS in the 5% dilated geometry is the most upstream. This is indicated in the WSS plot in figure 3(f): the restored vessels with varying dilation factors are reaching the minimum WSS extrema at staggered intervals along the dilated region, with the 15% dilated curve dipping the earliest the 5% curve dipping latest. It should also be noted that the observed minimum values of WSS in the dilated restoration geometries are all more extreme than the minimum WSS in the standard restoration geometry. In conclusion, this suggests that dilated geometries are more susceptible to CAD, as they feature a more extreme region of low WSS.

This observed WSS trend is reasonable. WSS is directly proportional to the gradient of the tangential component of velocity. Therefore, when increasing cross sectional area of the vessel lumen via dilation, velocity will decrease according to the continuity equation, resulting in low WSS.

### 3.3 Vorticity Observed in Restored LCX

An interesting characteristic is the presence of high vorticity in the FDR, as portrayed in figure 4. It is observed that a local maximum vorticity of 1266.2 s^−1^ along the vessel lies in the FDR of the undilated restored LCX, shown infigure 4(a). The local maximum is also illustrated in 4(e), where the peak in the graph represents the local maximum vorticity in the FDR. Similar to the undilated restoration geometry, the dilated vessels also contain a region of high vorticity near the formerly diseased region, as shown in figures 4(b,c,d). It is observed that the maximum vorticity in the FDR of the 5%, 10%, and 15% dilated vessels are 1035.38 s^−1^, 1164.27 s^−1^ and 1386.51 s^−1^ respectively. In this manner, the dilated vessels also exhibit an absolute maximum in vorticity in the same manner as the standard restored vessel, further validating vorticity as a flow characteristic associated with CAD in the LCX. The presence of vorticity can be linked to the curvature of the vessel, due to the fact that vorticity is dependent on radius of curvature. Therefore, both flow vorticity and vessel curvature could potentially be used as a metric to gauge risk for CAD development. To better establish these ideas, the relationship between vorticity and CAD can be further investigated by repeating studying more data.

**Figure 4:**
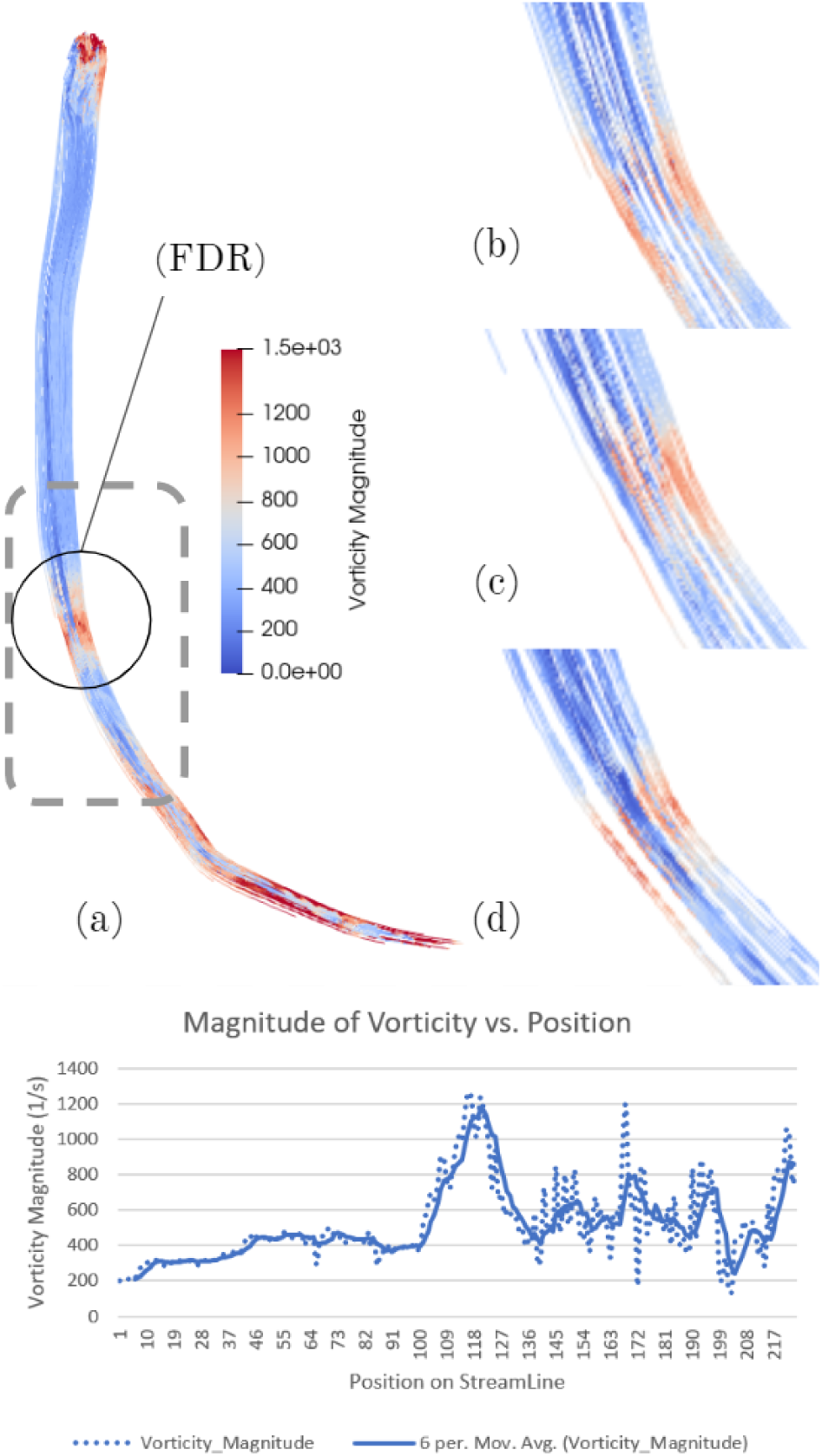
Vorticity in the restored LCX and dilated variations, s^−1^. (**a**) Vorticity in the undilated restored LCX. The FDR is indicated by a black circle. A band of red and orange shades is present in the FDR, implying high vorticity. (**b, c, d**) Vorticity in the 5%, 10%, and 15% dilated variations of the restored LCX respectively. Each plot is cropped such that it contains only the FDR. As shown, all three dilated variations feature the high vorticity characteristic in the FDR, indicated by the bands of red. (**e**) Magnitude of vorticity in the undilated restored LCX. All four restored vessels (1 undilated, 3 dilated) were analyzed for vorticity along its longitudinal axis, and the results of the undilated vessel are shown. A streamline passing through the vessel was generated, and vorticity values along the streamline spanning across the dashed gray rectangle labelled in (a) was graphed. In this manner, plots of vorticity vs. local element position were created, and a moving average of the vorticity values was also taken. The plot illustrates that on average, a local maximum of vorticity is present in the FDR. Generated plots for the dilated restored vessels are provided in the supplemental materials section.

## 4 Discussion

Recent advancements in cardiovascular simulation software have enabled patient-specific simulation for individualized diagnosis and treatment planning. This is particularly valuable considering that the role of hemodynamics in any disease scenario is highly individualized [Updegrove et al., 2017]. Tools such as Sim-Vascular have been key in research on individualized medicine, surgical planning and medical device design [Updegrove et al., 2017]. Recently, CTA derived fractional flow reserve became the first and only simulation-based technology approved by the United States Food and Drug Administration for CAD diagnosis; this method is being implemented by HeartFlow Inc, in Redwood City, California [Nørgaard et al., 2019]. Other noninvasive methods based on machine learning and reduced order models have also been increasing in popularity [Douglas et al., 2015]. Such methods can be improved as further analysis of the hemodynamics associated with CAD is done.

In this study, an analysis of the hemodynamics in a diseased and restored LCX was conducted. Through this comparison, a pressure drop across the stenosis in the diseased LCX was observed, which contrasted the smooth change in pressure across the restored LCX. This hints at the possibility of identifying sudden pressure drops in order to detect stenosis. Furthermore, it was observed that the region in which CAD developed was initially a region featuring low WSS and high vorticity. The low WSS characteristic of the restored geometry provides general validation for the restoration process, as this is an expected characteristic of a region that developed CAD. This in turn suggests that artificially restoring diseased vessel geometries to its initially healthy state is a promising method for investigating hemodynamics associated with the development of CAD. The vorticity characteristic is of interest as it may be associated with the curvature of the vessel. It has been shown in other studies that lesser curvature regions are atheroprone, and feature disturbed blood flow [see Heo et al., 2014, Chiu and Chien, 2011, Nigro et al., 2011]. Risk for the development of CAD in these regions can potentially be quantified using vorticity. Vorticity is defined as the curl of velocity: 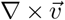. In natural coordinates:

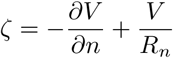

The first term is the vorticity resulting from shear, and the second term is the vorticity resulting from curvature of the path. As shown, vorticity is related to radius of curvature (*R*_*n*_). Hence, it is possible that vorticity may be used as a way to quantitatively analyze the risk of CAD development in regions with curvature. The procedure outlined in this study can be reused on more data to gain a generalized understanding and reinforce the relationship between vorticity and CAD in regions with curvature.

The initial simulation of the undilated restored LCX was validated by conducting simulations on dilated variations, where the artificially restored segments were dilated by factors of 5%, 10%, and 15%. This was done to test the tolerance of the patterns identified earlier, given that the standard, undilated restoration geometry may not accurately capture the true healthy geometry of the patient. The geometries with larger dilation factors featured a more extreme minimum WSS in the formerly diseased region. This implies that dilated vessels are more susceptible to the development of CAD. Similarly, the high vorticity characteristic identified in the undilated restored LCX was upheld in the dilated variations. It is likely that the vorticity contributes to the disturbed blood flow in the formerly diseased region, which has been identified as an atheroprone trait in other studies.

Interestingly, the research done by Li et al. [2018] also reached very similar conclusions regarding the hemodynamics associated with atherosclerotic stenosis. In that study, low WSS, low velocity, and apparent vortex was observed in the formerly diseased region of the repaired carotid vessel geometry. These findings corroborate with the observed low WSS and high vorticity in this study, providing further validation for the identified flow characteristics that are associated with CAD. However, validation for the application of artificial reconstruction to study the hemodynamics of diseased vessels is still needed to confirm the accuracy of the observed patterns.

The findings of this study and others can be advanced and reinforced by analyzing more data; however, this can only be accomplished with increased availability in public datasets. As aforementioned, the difficulties of obtaining the necessary data for analyzing the hemodynamics associated with CAD significantly limits this field of research. Steps have been taken in recent years to improve upon this: the National Institute of Health has recently released several large datasets, including the DeepLesion dataset created by Yan et al. [2018] which features over 32,000 CT images with a variety of lesions, accounting for 4,400 unique patients. This dataset was released with the intention of improving detection accuracy of lesions. While the DeepLesion dataset has improved the situation with public access to data in this field, it would be beneficial if a more focused dataset with CTA imaging of CAD patients were also made available. As further advancements in publicly-available datasets are made, CAD research, as well as the broader field of universal lesion detection can be advanced via machine learning methods [Yan et al., 2018].

To enable a more comprehensive analysis of the hemodynamics associated with CAD, the procedure used in this study can be modified, assuming the necessary data is available. First, it would be invaluable to analyze data from a large number of CAD patients for validation and more thorough pattern identification. It has also been shown that including side branches of the vessel in question improves the accuracy of WSS and pressure gradient calculations [Vardhan et al., 2019]. Hence, a potential improvement would be to include either the distal end of the LCX along with the second obtuse marginal artery or the entire coronary tree as part of the simulation process. Another potential improvement would be to implement pulsatile flow and a lumped parameter Windkessel model for the outlet boundary condition to more accurately represent blood flow through the coronary tree. Similarly, the elasticity of the vessel walls can be better represented by treating it as being deformable with uniform wall properties (Poisson’s ratio, elastic modulus, etc).

The accuracy of the segmentations can also be improved by not resorting to circular approximations of the vessel lumen, which may in turn influence the hemodynamic characteristics of the blood flow. Accurate segmentations can be achieved only via higher quality imaging, such as an optical coherence tomography. In addition, if the vessel segmentations are not approximated as circles, the restoration technique used in this study (a linear radius function of position) will no longer be valid. Hence, Voronoi diagram based reconstruction or mesh sweep reconstruction will be necessary. The effects of these techniques on the hemodynamics that are observed can help validate the patterns identified in this study.

More research is necessary to better understand the relationships between vorticity, WSS, curvature, and the development of CAD. A potential avenue would be to study the fluid dynamics of flows in artificially generated vessels with varying curvature to search for vorticity and other more established characteristics of atheroprone regions. Such simulations could help validate the findings in this study.

More fundamentally, rigorous biological explanations to these observed patterns are needed to confirm the relevance of the dynamics of blood flow and the development of CAD. Recent studies have involved protein kinase c zeta (PKC*ζ*) activity and its role in atherosclerosis and eventually CAD [Nigro et al., 2011]. As the understanding of hemodynamic factors and their contributions to CAD become clearer, risk analysis, diagnosis and treatment for CAD will become more accurate and effective.

